# Mitochondrial genome sequencing of marine leukemias reveals cancer contagion between clam species in the Seas of Southern Europe

**DOI:** 10.1101/2021.03.10.434714

**Authors:** Daniel Garcia-Souto, Alicia L. Bruzos, Seila Diaz, Sara Rocha, Ana Pequeño, Camila F. Roman-Lewis, Juana Alonso, Rosana Rodriguez, Damian Costas, Jorge Rodriguez-Castro, Antonio Villanueva, Luis Silva, Jose Maria Valencia, Giovanni Annona, Andrea Tarallo, Fernando Ricardo, Ana Bratos Cetinic, David Posada, Juan Jose Pasantes, Jose M. C. Tubio

**Author notes:** Denotes equal contribution (Daniel Garcia-Souto, Alicia L. Bruzos, Seila Diaz). Correspondence to: Dr Jose M. C. Tubio, Genomes and Disease, Molecular Medicine and Chronic Diseases Research Centre (CIMUS), University of Santiago de Compostela, Santiago de Compostela 15706, Spain, Dr. Daniel Garcia-Souto, Genomes and Disease, Molecular Medicine and Chronic Diseases Research Centre (CIMUS), University of Santiago de Compostela, Santiago de Compostela 15706, Spain.

## Abstract

Clonally transmissible cancers are tumour lineages that are transmitted between individuals via the transfer of living cancer cells. In marine bivalves, leukemia-like transmissible cancers, called hemic neoplasia, have demonstrated the ability to infect individuals from different species. We performed whole-genome sequencing in eight warty venus clams that were diagnosed with hemic neoplasia, from two sampling points located more than 1,000 nautical miles away in the Atlantic Ocean and the Mediterranean Sea Coasts of Spain. Mitochondrial genome sequencing analysis in tumour tissues from neoplastic animals revealed the coexistence of haplotypes from two different clam species. Phylogenies estimated from mitochondrial and nuclear markers confirmed this leukemia originated in striped venus clams and later transmitted to clams of the species warty venus, in which it survives as a contagious cancer. The analysis of mitochondrial and nuclear gene sequences supports all studied tumours belong to a single neoplastic lineage that spreads in the Seas of Southern Europe.

## INTRODUCTION

Cancers are clonal cell lineages that arise due to somatic changes that promote cell proliferation and survival (Stratton, Campbell, & Futreal, 2009). Although natural selection operating on cancers favours the outgrowth of malignant clones with replicative immortality, the continued survival of a cancer is generally restricted by the lifespan of its host. However, clonally transmissible cancers – from now on, transmissible cancers – are somatic cell lineages that are transmitted between individuals via the transfer of living cancer cells, meaning that they can survive beyond the death of the individual that spawned them (Murchison, 2008). Naturally-occurring transmissible cancers have been identified in dogs (Murchison et al., 2014), Tasmanian devils (Murchison et al., 2012) and, more recently, in marine bivalves (Metzger, Reinisch, Sherry, & Goff, 2015; Metzger et al., 2016).

Hemic neoplasia (HN), also called disseminated neoplasia, is a type of leukemia cancer found in multiple species of bivalves, including oysters, mussels, cockles and clams (Carballal, Barber, Iglesias, & Villalba, 2015). Although these leukemias represent different diseases across bivalve species, they have been classically grouped under the same term because neoplastic cells share morphological features (Carballal et al., 2015). HN has a clonal transmissible behaviour (Metzger et al., 2015), in which the haemocytes (i.e., the cells that populate the haemolymph and play a role in the immune response) are likely to be transmitted through marine water. In late stages of the disease, leukemic cells invade the surrounding tissues and animals die because of the infection (Carballal et al., 2015). Despite leukemic haemocytes are typically transmitted between individuals from the same species, on occasion they can infect and propagate across populations from a second, different bivalve species (Metzger et al., 2016; Yonemitsu et al., 2019), representing interesting models for the understanding of the genetic causes of cancer transmissibility and metastasis.

Marine transmissible cancers likely move using ocean currents to colonize new regions, although they may also be unintentionally introduced by the action of man in disease-free regions (Yonemitsu et al., 2019). Hence, these cancers represent a potential threat for the ecology of the marine environment, which claims for the necessity of their identification and genetic characterization for their monitoring and prevention. Here, we use next-generation sequencing to identify a new marine leukemia that is transmitted between two different clam species in the Atlantic Ocean and the Mediterranean Sea of Southern Europe: warty venus (*Venus verrucosa*) and striped venus (*Chamelea gallina*) clams.

## RESULTS AND DISCUSSION

We investigated the prevalence of hemic neoplasia in the warty venus clam (*V. verrucosa*), a saltwater bivalve found in the Atlantic Coast of Europe and the Mediterranean Sea. We collected 345 clam specimens from six sampling regions in the Atlantic and the Mediterranean coasts of Europe across five different countries, including Spain, Portugal, France, Ireland and Croatia (**Figure 1a**; **Supplementary Table 1**). Cyto-histological examination identified hemic neoplasia-like tumours in eight specimens from two sampling points in Spain. Three HN-positive specimens (ERVV17-2995, ERVV17-2997, ERVV17-3193) were collected in Galicia, northwest of the Iberian Peninsula in the Atlantic Ocean, and another five specimens (EMVV18-376, EMVV18-391, EMVV18-395, EMVV18-400) were collected in the Balearic Islands, bathed by the Mediterranean Sea (**Figure 1a**). Four of these specimens (ERVV17-2995, ERVV17-3193, EMVV18-391, EMVV18-395) showed a severe form of the disease – classified as N3 stage –, which is characterized by high levels of neoplastic cells in the haemolymph and infiltration of neoplasia in the surrounding tissues **(Figure 1b-c, Supplementary Figure 1)**; three specimens (ERVV17-2997, EMVV18-373, EMVV18-376) were diagnosed with a light form of the disease – N1 stage –, characterized by low levels of neoplastic cells in the haemolymph and no infiltration of tissues **(Supplementary Figure 1)**; and one specimen (EMVV18-400) was found that was affected with an intermediate form of the disease – N2 stage –, characterized by intermediate level of neoplastic cells in the haemolymph and low infiltration of tissues **(Supplementary Figure 1)**. Electron microscopy analysis through gill’s ultrathin sections from two neoplastic warty venus specimens revealed tumour cells with a round shape and a pleomorphic nucleus, which are morphological features that generally characterize bivalves’ hemic neoplasia, although warty venus tumour cells also show the ability to emit pseudopodia (**Figure 1d**). In private sample collections from our collaborators, we found one additional neoplastic warty venus specimen (EVVV11-02) from Galicia, which was collected in 2011. This animal showed abnormal karyotypes in the gills, consisting of increased chromosome numbers, abnormal chromosome morphologies and location of cytogenetic markers, which was suggestive of hemic neoplasia (**Figure 1e**).

**Table 1.** Clam specimens and tissues sequenced with Illumina paired-ends. Sixteen specimens (eight neoplastic and eight non-neoplastic) from three different clam species (*V. verrucosa, C. gallina, C. striatula*) were sequenced with Illumina paired-ends. Columns 5 and 6 show the number of reads generated for the host tissue (when neoplastic, matched-normal tissue was foot) and the tumoral haemolymph, respectively. (*) denotes the only available tissue from this neoplastic animal, collected in 2011, were gills. (#) denotes hemic neoplasia stage was not determined because cyto-histological examination was not possible in this individual, which was diagnosed by cytogenetics.

| Clam species | Specimen origin | Specimen code | Diagnosis | Host reads | Tumoral haemolymph reads |
| --- | --- | --- | --- | --- | --- |
| *V. verrucosa* | Galicia, Spain | ERVV17-2995 | N3 | 833M | 919M |
| *V. verrucosa* | Galicia, Spain | ERVV17-2997 | N1 | 766M | 598M |
| *V. verrucosa* | Galicia, Spain | ERVV17-3193 | N3 | 739M | 850M |
| *V. verrucosa* | Balearic Islands, Spain | EMVV18-376 | N1 | 784M | 849M |
| *V. verrucosa* | Balearic<br>Islands, Spain | EMVV18-391 | N3 | 617M | 623M |
| *V. verrucosa* | Balearic<br>Islands, Spain | EMVV18-395 | N3 | 697M | 679M |
| *V. verrucosa* | Balearic<br>Islands, Spain | EMVV18-400 | N1 | 782M | 1133M |
| *V. verrucosa* | Galicia, Spain | EVVV11-02 | N<sup>#</sup> | 743M* | -* |
| *V. verrucosa* | Dubrovnik,<br>Croatia | CSVV18-1052 | Healthy | 161M | - |
| *V. verrucosa* | Mahon, Spain | EMVV18-385 | Healthy | 143M | - |
| *V. verrucosa* | Granville,<br>France | FGVV18-183 | Healthy | 752M | - |
| *V. verrucosa* | Galway,<br>Ireland | IGVV19-666 | Healthy | 155M | - |
| *V. verrucosa* | Lisbon,<br>Portugal | PLVV18-2249 | Healthy | 163M | - |
| *C. gallina* | S. Benedetto,<br>Italy | IMCG15-69 | Healthy | 147M | - |
| *C. gallina* | Cadiz, Spain | ECCG15-201 | Healthy | 752M | - |
| *C. striatula* | Vigo, Spain | EVCS14-09 | Healthy | 706M | - |

**Figure 1.**
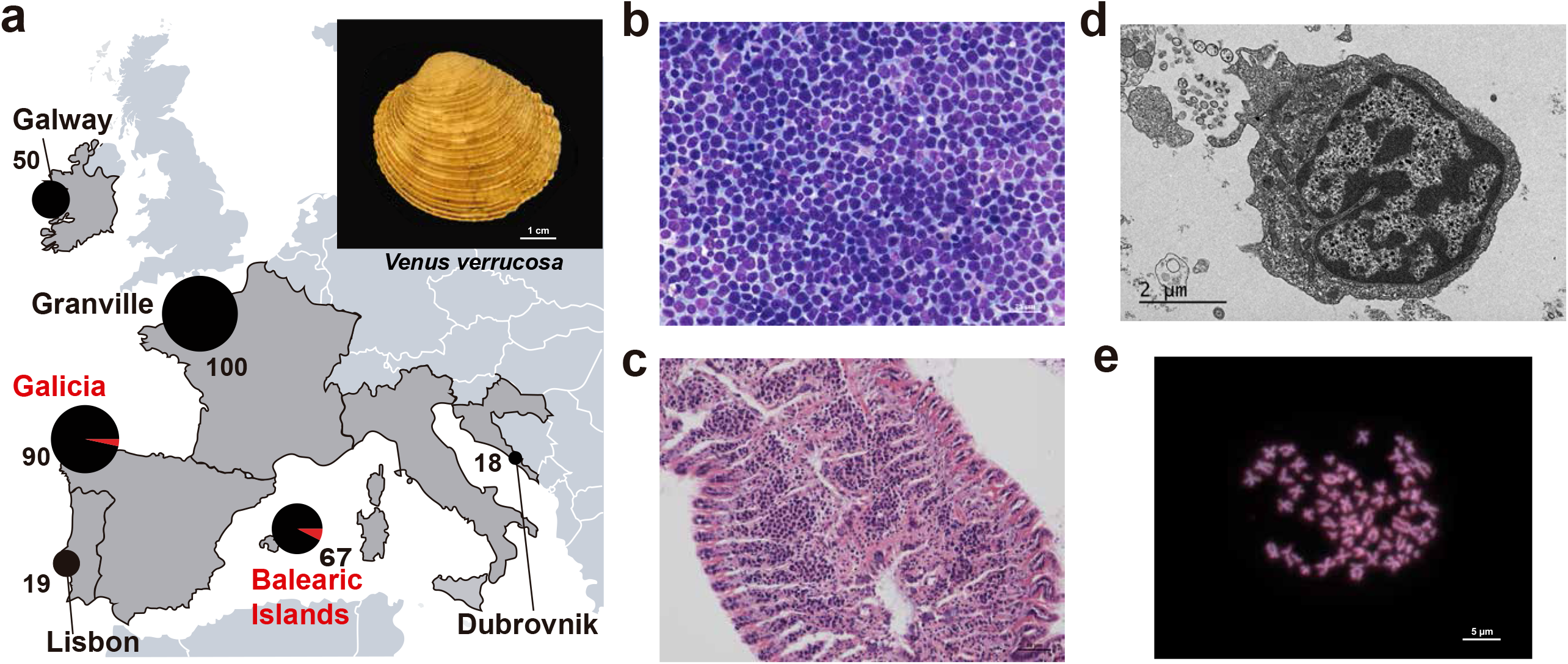
Geographical location of warty venus (*V. verrucosa*) specimens and diagnosis of hemic neoplasia. (a) Locations of *V. verrucosa* clams collected and specimens diagnosed with hemic neoplasia. Sizes of the pie charts are in proportion with the number of samples collected (number of samples is shown next to each pie chart). Pie charts show the proportion of samples with hemic neoplasia (black, no neoplastic specimens; red, neoplastic specimens). Top-right corner shows a representative specimen of the species *V. verrucosa*. (b) Cytological examination of haemolymph smear (Hemacolor stain) from a remarkable example of a *V. verrucosa* specimen with high-grade (N3 stage) hemic neoplasia showing neoplastic cells that replaced normal haemocytes. (c) From the same N3 specimen above, hematoxylin and eosin-stained light micrograph of histological section from the gills shows vessels filled with neoplastic cells. (d) Transmission electron microscopy analysis of a *V. verrucosa* hemic neoplasia tumour cell shows a round shape, pseudopodia “p”, pleomorphic nucleus “n” with scattered heterochromatin, and mitochondria “m”. (e) Metaphase chromosomes from a neoplastic cell from infiltrated gills of *V. verrucosa* specimen EVVV11-02 reveals abnormal chromosome number (>19 pairs), morphology and cytogenetic markers. Chromosomes stained with DAPI and PI. Scale bar = 5 μm.

To illuminate some biological insights into the clonal dynamics of this cancer, we carried out whole-genome sequencing with Illumina paired-ends in DNA samples isolated from the tumoral haemolymph from eight out of the nine neoplastic specimens (**Table 1**). Their feet were also sequenced, as foot typically represents the tissue with lower infiltration of neoplastic cells, making it a good candidate tissue to act as “matched-normal” (i.e., host tissue). As for the animal with an abnormal karyotype (EVVV11-02) that was compatible with hemic neoplasia, we sequenced the only tissue available, which were gills (**Table 1**). Only one neoplastic specimen (EMVV18-373) that had very low proportion of tumour cells in its haemolymph was excluded from the sequencing. Then, we mapped the paired-end reads onto a dataset containing non-redundant mitochondrial Cytochrome C Oxidase subunit 1 (*Cox1*) gene references from 137 Vererid clam species. In six out of eight sequenced neoplastic specimens, the results revealed an overrepresentation (>99%) of reads in the sequenced tissues mapping *Cox1* DNA sequences that exclusively identified two different clam species (**Figure 2a**): the expected one, warty venus clam (*V. verrucosa*), and a second, unexpected one, the striped venus (*C. gallina*), a clam that inhabits the Mediterranean Sea (**Figure 2b**). Preliminary analysis by PCR and capillary sequencing of *Cox1* in the haemolymph of two neoplastic specimens, EMVV18-373 and EVVV11-02, revealed an electropherogram with overlapping peaks apparently containing two different haplotypes that match the reference *Cox1* sequences for warty venus and striped venus (**Figure 2c**).

**Figure 2.**
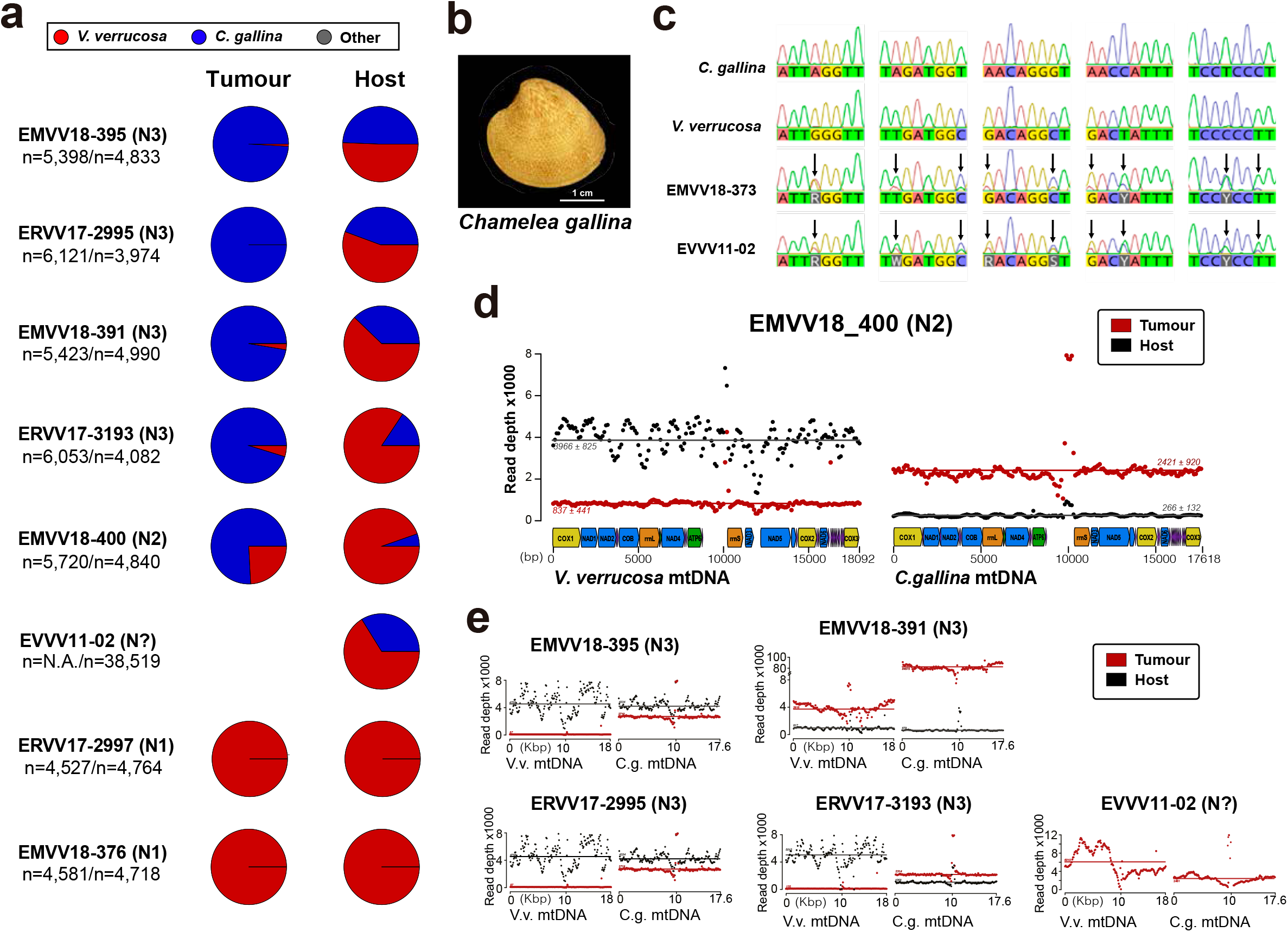
Mitochondrial DNA sequencing analyses reveal cancer contagion between warty venus (*V. verrucosa*) and striped venus (*C. gallina*) clam species. (a) In eight warty venus specimens sequenced with Illumina paired-ends, the pie charts show the proportion of reads mapping *Cox1* reference sequences from 137 different *Verenidae* species, including *V. verrucosa* (red), *C. gallina* (blue) and the remaining species (grey). Two different tissues were sequenced: the tumour tissue (left pie chart), typically haemolymph, and the host/matched-normal tissue (right pie chart), typically foot. Note that for specimen EVVV11-02 only the host/matched-normal tissue (gills) was available. “n” denotes the total number of reads mapping the *Cox1* reference for the tumour tissue (left), and the host tissue (right). (b) Representative specimen of the species *C. gallina*. (c) Capillary sequencing electropherograms of mitochondrial *Cox1* gene fragments from two neoplastic *V. verrucosa* specimens (EMVV18-373 and EVVV11-02) and two healthy reference specimens from *V. verrucosa* and *C. gallina*. The results show overlapping peaks (arrows) in the sequenced tissues from the neoplastic animals, which suggest coexistence of mtDNA haplotypes from two clam species. (d) In *V. verrucosa* neoplastic (N2-stage) specimen EMVV18-400, mtDNA read depth shows different proportion of warty venus and striped venus mtDNA haplotypes in the tumour tissue (haemolymph) and the matched-normal tissue (foot). (e) Similar patterns were found in the remaining five *V. verrucosa* neoplastic specimens that were sequenced.

These results suggested cancer contagion between the two clam species of the family Veneridae. Hence, we employed bioinformatic tools to reconstruct the full mitochondrial DNA (mtDNA) genomes in representative animals from the two species involved. The method provided two sequences, 18,092 bp and 17,618 bp long, for *V. verrucosa* and *C. gallina*, respectively (**Supplementary Figure 2**). Then, we mapped the paired-end sequencing data from the six neoplastic specimens with evidence of interspecies cancer transmission onto the two reconstructed species-specific mtDNA genomes. This approach confirmed the coexistence of two different mtDNA haplotypes in the examined neoplastic samples, matching the canonical mtDNA genomes from the two clam species. For example, in a remarkable N2-stage specimen (EMVV18-400), this analysis revealed different proportion of tumour and host warty venus and striped venus mtDNA molecules in the two tissue types sequenced, being the striped venus mtDNA the most abundant in the tumour tissue (i.e., haemolymph), where tumour cells are the dominant cell type, and the lower in the matched-normal tissue (i.e., infiltrated foot), where tumour cells represent a minor fraction of the total (**Figure 2d**). Similar results were obtained for the remaining five neoplastic individuals (**Figure 2e)**.

To further investigate the evolutionary origins of this cancer, we sequenced with Illumina paired-ends an additional set of eight healthy (i.e., non-neoplastic) clams from three different *Veneridae* species, including five more warty venus specimens (EMVV18-385, IGVV19-666, FGVV18-183, CSVV18-1052, PLVV18-2249) from five different countries, two striped venus specimens (IMCG15-69, ECCG15-201) from two countries, and one specimen (EVCS14-09) from its sibling species *Chamelea striatula*, a type of striped venus clam that inhabits the Atlantic Ocean from Norway to the Gulf of Cadiz in Spain. This made a total of 16 Veneridae specimens sequenced, all listed in **Table 1** (**Supplementary Table 1**). We reconstructed the reference mtDNA genome of the third species, *C. striatula*, as we did for *V. verrucosa* and *C. gallina*. Then, the sequencing reads from the 16 animals were mapped onto the three species-specific reference mitochondrial genomes. A consensus mtDNA sequence was inferred from the mapping data for each one of the 16 clam specimens sequenced; an approach that in the neoplastic animals reconstructed two different haplotypes due to the presence of DNA from two different species. Then, we performed a preliminary phylogenetic analysis based on the alignment of mtDNA sequences from 13 coding and two RNA gene sequences, altogether encompassing ~14 kb. The results show that tumour and non-tumour sequences from neoplastic warty venus specimens split out in the phylogenetic tree conforming two different clusters, and that tumoral warty venus sequences clustered all together and closer to striped venus than to warty venus specimens (**Figure 3a**). This data supports the existence of a single cancer clone originated in the striped venus clam *C. gallina* that was transmitted to *V. verrucosa*.

**Figure 3.**
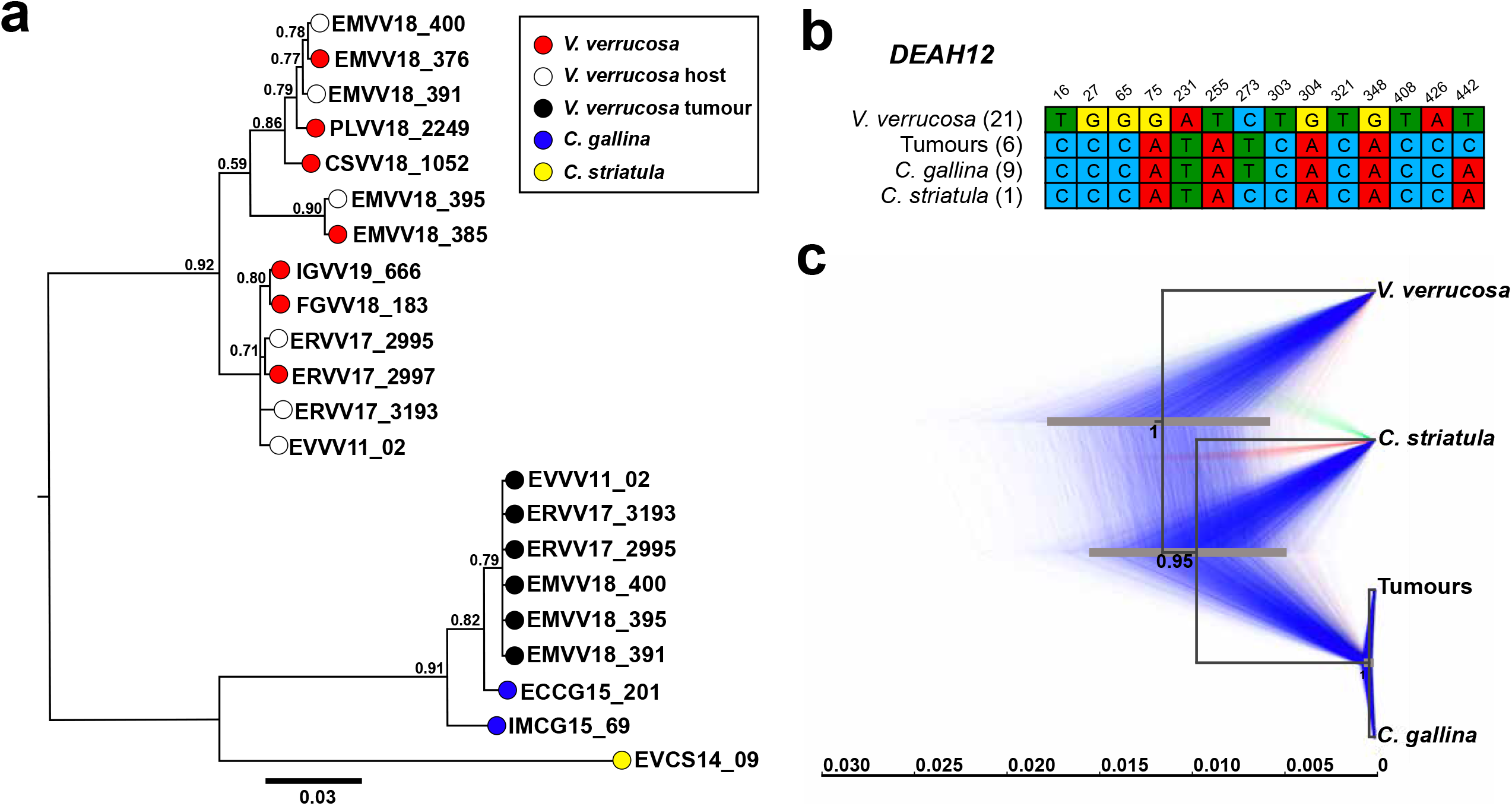
Phylogenetic analyses confirm a single cancer lineage spreading in populations of the warty venus (*V. verrucosa*) that was originated in the striped venus (*C. gallina*). (a) Molecular phylogeny using Bayesian inference inferred on the alignment of all mitochondria coding genes and rRNA gene sequences (15 loci) that includes six neoplastic *V. verrucosa* specimens with evidence of cancer contagion from *C. gallina*. Bootstrap values are shown above the branches. (b) Single-nucleotide variants discriminating between *V. verrucosa* hosts and *V. verrucosa* tumours along a 441 bp region of nuclear gene *DEAH12*. (c) Bayesian phylogeny of *V. verrucosa*, their tumours and *Chamelea sp*. based on the entire mitochondrial DNA and nuclear marker *DEAH12*. A Maximum Clade Credibility tree (MCC) is shown, with posterior probabilities below the branches, and 95% highest probability density (HPD) intervals of node heights as grey bars. A tree cloud including 1,000 trees represents the range of alternative topologies, in which blue is the most common set of topologies, red the second most common one, and green the remaining.

Transmissible cancers are known to occasionally acquire mitochondria from transient hosts (Strakova et al., 2016; Strakova et al., 2020), which can lead to misinterpretation of their evolutionary history. Thus, we looked for nuclear markers to confirm the striped venus origin of this cancer lineage. We performed a preliminary draft assembly of the warty venus and the striped venus nuclear reference genomes, using the paired-end sequencing data from two non-neoplastic animals. Then, we used bioinformatic approaches to identify single copy nuclear genes that were homologous between the two species. From this analysis we selected a 2.9 kb-long region from *DEAH12*, a gene that encodes for an ATP-dependent RNA helicase. With the idea of identifying differentially fixed single-nucleotide variants (SNVs) between both species, we performed PCR amplification and capillary sequencing on a 441 bp fragment from *DEAH12*, on 12 non-neoplastic warty venus and 9 striped venus specimens. In this way, we identified 28 SNVs from 14 nucleotide positions that allow to discriminate between the two species (**Fig. 3b**). This set of variants was later employed to filter the Illumina short reads from the alignments of warty venus neoplastic specimens matching either warty venus or striped venus genotypes, and to obtain the consensus sequences from tumour and non-affected tissue in each sequenced specimen. We obtained the *DEAH12* reference sequence for *C. striatula*, and these sequences were used to construct a multilocus species tree based on the alignment of the mitochondrial DNA and the nuclear gene from healthy/neoplastic *V. verrucosa, C. gallina*, and *C. striatula*. The phylogeny confirmed a single clade of warty venus tumours that is closer to striped venus specimens than to non-neoplastic warty venus sequences from the same diseased specimens, while the non-neoplastic sequences conformed a more distant warty venus lineage (**Figure 3c**). To find out whether this cancer is alive in the clam species where it first arose, we performed a screening for its presence in natural populations of striped venus clams from the species *C. gallina* and *C. striatula* at five additional sampling points across two countries (**Supplementary Table 1**), including Spain (n=109) and Italy (n=57). Histological analysis did not show any traces of hemic neoplasia in these specimens. The virtual absence of this tumour in natural populations of striped venus clams may suggest that today this leukemia is being mainly, if not exclusively, transmitted between specimens of the recipient species, warty venus. However, further sampling in other regions across the striped venus area of distribution may be necessary to confirm these findings.

The results provided here reveal the existence of a transmissible leukemia originated in a striped venus clam, most likely from the species *C. gallina*, which was transmitted to a second species, the warty venus clam (*V. verrucosa*), and among whose specimens it currently propagates. We identified this parasitic cancer in warty venus clams from two sampling points that are more than 1,000 nautical miles away in the coasts of Spain, bathed by two different seas, the Atlantic Ocean and the Mediterranean Sea. The analysis of mitochondrial and nuclear gene sequences revealed no nucleotide divergence between the seven tumours sequenced, which supports that all belong to the same neoplastic lineage that spreads between Veneridae clams in the Seas of Southern Europe. Although we ignore the age of this cancer clone, we can confirm it arose before 2011, when the neoplastic warty venus specimen EVVV11-02 was collected. The apparent lack of genetic variation between all tumours, even from distant sampling points, suggests either that this cancer is very recent, or that it may have been unintentionally scattered by the action of man, a way of transmission that has been proposed for other bivalve transmissible cancers (Yonemitsu et al., 2019).

## SUPPLEMENTARY FIGURES

**Supplementary Figure 1. Histological diagnosis of hemic neoplasia in six warty venus (V. verrucosa) specimens**. Hematoxylin and eosin-stained photomicrographs of gill, digestive (d), gonad (male (m) & female (f)) and foot of warty venus (*V verrucosa*) specimens diagnosed with different stages of hemic neoplasia: high (N3), medium (N2) and light (N1). In the N3 stage, neoplastic cells infiltrate the connective tissue and vessels of different organs (A-L), and show low infiltration of foot (C,F,I,L). In N2 stage, cell groups are observed in different organs such as gills (M) and are not detected in the foot (O). In N1 stage, groups of neoplastic or isolated cells are detected in gill sinuses (P, S, V). Arrows show isolated cells. Asterisks show group of neoplastic cells. Scale bar 200µm.

**Supplementary Figure 2. Draft reference mtDNA genome assemblies reconstructed for *V. verrucosa, C. gallina* and *C. striatula***. Preliminary reference mtDNA genome assembly obtained from paired-end sequencing data, together with gene annotation.

## SUPPLEMENTARY TABLES

**Supplementary Table 1. Sampling data of 570 specimens analysed in this study**.

## MATERIALS AND METHODS

### Sampling of clam specimens

We collected 570 clam specimens from three different species, from the following countries and locations (**Supplementary Table 1)**. *V. verrucosa* clams were collected in Spain (Galicia, n=90; Balearic Islands, n=67), France (Granville, n=100), Croatia (Dubrovnik, n=18), Portugal (Lisbon, n=19), and Ireland (Galway, n=50). *C. gallina* clams were collected in Spain (Cadiz, n=50; Mallorca, n=50), and Italy (Naples, n=50; Cattolica, n=57). *C. striatula* clams were collected in Spain (Combarro, n=9). Additionally, we recruited samples from the following specimens from private collections: one *V. verrucosa* clam collected in 2011 in Spain (Islas Cies), four *C. gallina* collected in 2015 in Italy (San Benedetto de Tronto), five *C. gallina* collected in 2015 in Spain (Huelva), and one *C. striatula* collected in 2014 in Spain (Marin).

### Diagnosis of hemic neoplasia

We followed standard cytological and/or histological protocols to diagnose hemic neoplasia in the clam specimens. However, only histological examination resulted decisive for the diagnosis, particularly in early stages of the disease. Briefly, for each animal, we extracted 300-2000 μl of haemolymph from the posterior adductor muscle using a 5 ml syringe with a 23G needle. The haemolymph (50 μl) was diluted in cold Alserver’s antiaggregant solution to a 1:4 concentration, and spotted by centrifugation (130g, 4ºC, 7 min) onto a microscope slide using cytology funnel sample chambers to produce a cell monolayer. Haemolymph smears were fixed and stained with Hemacolor solutions from Sigma-Aldrich and subsequently examined with a light microscope for the diagnosis of hemic neoplasia. Tissues (visceral mass, gills, mantle and foot) were dissected, fixed in Davidson’s solution and embedded in paraffin. Then, 5 μm thick sections from each tissue were microdissected and stained with Harris’ haematoxylin and eosin and examined using a light microscope for histopathological analysis. Hemic neoplasia was diagnosed and classified according to three disease stages (i.e., N1, N2 or N3) as follows. N1 stage: small groups of leukemic cells were detected only in the vessels of the gills and in the connective tissue surrounding the digestive tubules. N2 stage: leukemic cells spread to different organs, conforming small groups in the connective tissue that surrounds the digestive gland and the gonadal follicles, branchial sinuses and mantle. N3 stage: leukemic cells invade the filaments, completely deforming the plica structure in the gill, invade the connective tissue surrounding the gonadal follicles and the digestive gland; in the mantle, they invade the connective tissue, but in the muscle fibres of the mantle and foot, cells appear isolated or in small groups and in lower intensity than in other tissues.

### Electron microscopy analysis

Four *V. verrucosa* specimens (two non-neoplastic, ERVV17-2993 and ERVV17-2992, and two with high grade of hemic neoplasia, ERVV17-2995 and ERVV17-3193) were processed for transmission electron microscopy (TEM) as follows: 2 mm sections of gills and digestive glands were fixed in 2.5% glutaraldehyde seawater for 2 h at 4°C. Then, tissues were post-fixed in 1% osmium tetroxide in sodium cacodylate solution and embedded in Epon resin. Ultrathin sections were stained with uranyl acetate and lead citrate and examined in a JEM-1010 transmission electron microscope.

### Cytogenetics

The karyotype of a neoplastic *V. verrucosa* specimen (EVVV11-02) was obtained as follows. After colchicine treatment (0.005%, 10 h), gills were dissected, treated with a hypotonic solution, and fixed with ethanol and acetic acid. Small pieces of fixed gills were disaggregated with 60% acetic acid to obtain cell suspensions that were spread onto preheated slides. Chromosome preparations were stained with DAPI (0.14 μg/ml) and PI (0.07 μg/ml) for 8 minutes, mounted with antifade medium, and photographed.

### De-novo assembly of mitochondrial genomes and annotation

We performed whole-genome sequencing on 23 samples from 16 clam specimens, which includes eight neoplastic and eight non-neoplastic animals (**Table 1**), with Illumina paired-end libraries of 350 bp insert size and reads 150 bp long. Then, we run MITObim v1.9.1 (Hahn, Bachmann, & Chevreux, 2013) to assemble the full mitochondrial genome of all sequenced samples, using gene baits from the following *Cox1* and *16S* reference genes to prime the assembly of clam mitochondrial genomes: *V. verrucosa* (*Cox1*, with GenBank accession number KC429139, *16S*: C429301), *C. gallina* (*Cox1*: KY547757, *16S*: KY547777) and *C. striatula* (*Cox1*: KY547747, *16S*: KY547767). The draft sequences were polished twice with Pilon v1.23 (Walker et al., 2014), and conflictive repetitive fragments from the mitochondrial control region were resolved using long read sequencing with Oxford Nanopore technologies (ONT) on a set of representative samples from each species and tumours. ONT reads were assembled with Miniasm v0.3 (H. Li, 2016) and corrected using Racon v1.3.1 (Vaser, Sovic, Nagarajan, & Sikic, 2017). Protein-coding genes, rDNAs and tRNAs were annotated on the curated mitochondrial genomes using MITOS2 web server (Bernt et al., 2013), and manually curated to fit ORFs as predicted by ORF-FINDER (Rombel, Sykes, Rayner, & Johnston, 2002).

### PCR and capillary sequencing of *Cox1*

PCR primers were designed with Primer3 v2.3.7 (Koressaar et al., 2018) to amplify a fragment of 354 bp from the *Cox1* mitochondrial gene of *V. verrucosa* and *C. gallina* (F: CCT ATA ATA ATT GGK GGA TTT GG, R: CCT ATA ATA ATT GGK GGA TTT GG). PCR products were purified with ExoSAP-IT and sequenced by Sanger sequencing.

### Mitochondrial genome coverage analysis

We mapped the paired-end sequencing data from healthy and neoplastic tissues from all neoplastic samples onto the *V. verrucosa* and *C. gallina* reference mitochondrial genomes using BWA-mem v0.7.17-r1188 (H. Li & Durbin, 2009) with default parameters. Duplicate reads were marked with Picard 2.18.14 and removed from the analysis. Read coverage depth was computed with samtools v1.9 (H. Li et al., 2009), reduced by computing the average in windows of 100 bp and plotted with R v3.5.3.

### Draft assembly of nuclear reference genomes, identification of variable single-copy orthologous nuclear loci and SNPs

We run the megahit v1.1.3 assembler (D. Li, Liu, Luo, Sadakane, & Lam, 2015) on the Illumina paired-end sequencing data to obtain partial nuclear genome assemblies of *V. verrucosa, C. gallina* and *C. striatula*. Then, single copy genes were predicted with Busco v.3.0.2 (Seppey, Manni, & Zdobnov, 2019). Candidate genes were considered if they (1) were present in the genomes of the three species, and (2) showed variant allele frequencies (VAF) at exclusively 0, 0.5 or 1.0 in all the sequenced healthy (non-neoplastic) specimens. A single gene was selected by these criteria, a 3,914 bp long fragment of *DEAH12*, a gene encoding for an ATP-dependent RNA helicase. PCR primers were designed with Primer3 v2.3.7 (Koressaar et al., 2018) to amplify and sequence a 441 bp region of the *DEAH12* nuclear gene on specimens from the species *V. verrucosa* and *C. gallina*; F: AGG TAT GCT GAA ACA AAC ACT T, R: ACG ACA AAT TTG ATA CCT GGA AT. Then, PCR amplification and Sanger sequencing of this region were performed in non-neoplastic animals from both species (11 *V. verrucosa* and nine *C. gallina*). We screened for differentially fixed single-nucleotide variants (SNVs) between both species using functions in the R package Exploratory Analysis of Genetic and Genomic Data *adegenet* (Jombart & Ahmed, 2011). These SNPs were later employed to filter the Illumina short reads matching either *V. verrucosa* or *C. gallina* genotypes from neoplastic animals, and to obtain the consensus sequences from tumour and non-affected tissue in each sequenced specimen.

### Phylogenetic analyses

Mitochondrial sequences for 13 coding genes and two rDNA genes from 16 specimens (six neoplastic, 10 non-neoplastic) were extracted from the paired-end sequencing data, concatenated, and subjected to multiple alignment with MUSCLE v3.8.425 (Edgar, 2004). The best-fit model of nucleotide substitution for each individual gene was selected using JModelTest2 (Darriba, Taboada, Doallo, & Posada, 2012) and a partitioned Bayesian reconstruction of the phylogeny was performed with MrBayes v3.2.6 (Ronquist et al., 2012). Two independent Metropolis-coupled Markov Chain Monte Carlo (MCMC) analyses with four chains in each were performed. Each chain was run for 10 million generations, sampling trees every 1,000 generations. Convergence between runs was assessed using Tracer (Rambaut, Drummond, Xie, Baele, & Suchard, 2018).

*DEAH12* sequences from 31 specimens (six neoplastic, 25 non-neoplastic) were extracted from the paired-end sequencing data, and subjected to multiple alignment using MUSCLE v3.8.425 (Edgar, 2004). Then, a “species tree” was inferred with the starBEAST multispecies coalescent (MSC) model, as implemented in BEAST v2.6.2 (R. Bouckaert et al., 2019). This analysis was performed using a Yule speciation prior and strict clock, with the best-fit model of nucleotide substitution obtained with jModelTest2 (Darriba et al., 2012) on both the concatenated mitochondrial haplotypes (13 protein-coding and two rRNAs genes) and unphased data from *DEAH12* nuclear marker. The four mitochondrial groups observed on the mitogenome analysis (*V. verrucosa, C. gallina, C. striatula* and Tumor) were defined as tips for the species tree. A single MCMC of 10 million iterations, with sampling every 1000 steps, was run. A burn-in of 10%, was implemented to obtain ESS values above 200 with TRACER v1.7.1 (Rambaut et al., 2018) and the resulting posterior distributions of trees were checked with DENSITREE v2.1 (R. R. Bouckaert, 2010). A maximum clade credibility (MCC) tree was obtained with TreeAnotator to summarize information on topology, with 10% burn-in and Common Ancestors for the node heights.

## ACKNOWLEDGEMENTS

We thank the Galicia Supercomputing Centre (CESGA) for the availability of informatic resources.

## ADDITIONAL INFORMATION

### Data availability

Nucleotide data for the mitochondrial DNA assemblies has been uploaded to GenBank under accession numbers MW662590-MW662611 and will be released upon publication or request. These correspond to reference healthy animals of *Venus verrucosa* (MW662590, MW662593, MW662607, MW662608 and MW662610), *Chamelea gallina* (MW662591 and MW662609), *Chamelea striatula* (MW662611) and to both normal (MW662592, MW662595, MW662597, MW662599, MW662601, MW662602, MW662604, MW662606) and tumoral (MW662594, MW662596, MW662598, MW662600, MW662603, MW662605) mitochondrial DNAs of neoplastic animals.

Genomic data (Illumina paired-end sequencing) from the mitogenomes analysed in this manuscript are allocated in DRYAD and available through the following link: https://datadryad.org/stash/share/eSwRiQ_aSTTCJMWo7bFrPp6FQVDViqkrsV1td6RHWEg

