## Supplementary figures and images for "Mitochondrial genome sequencing of marine leukemias reveals cancer contagion between clam species in the Seas of Southern Europe"

### Supplementary Figures 1-2

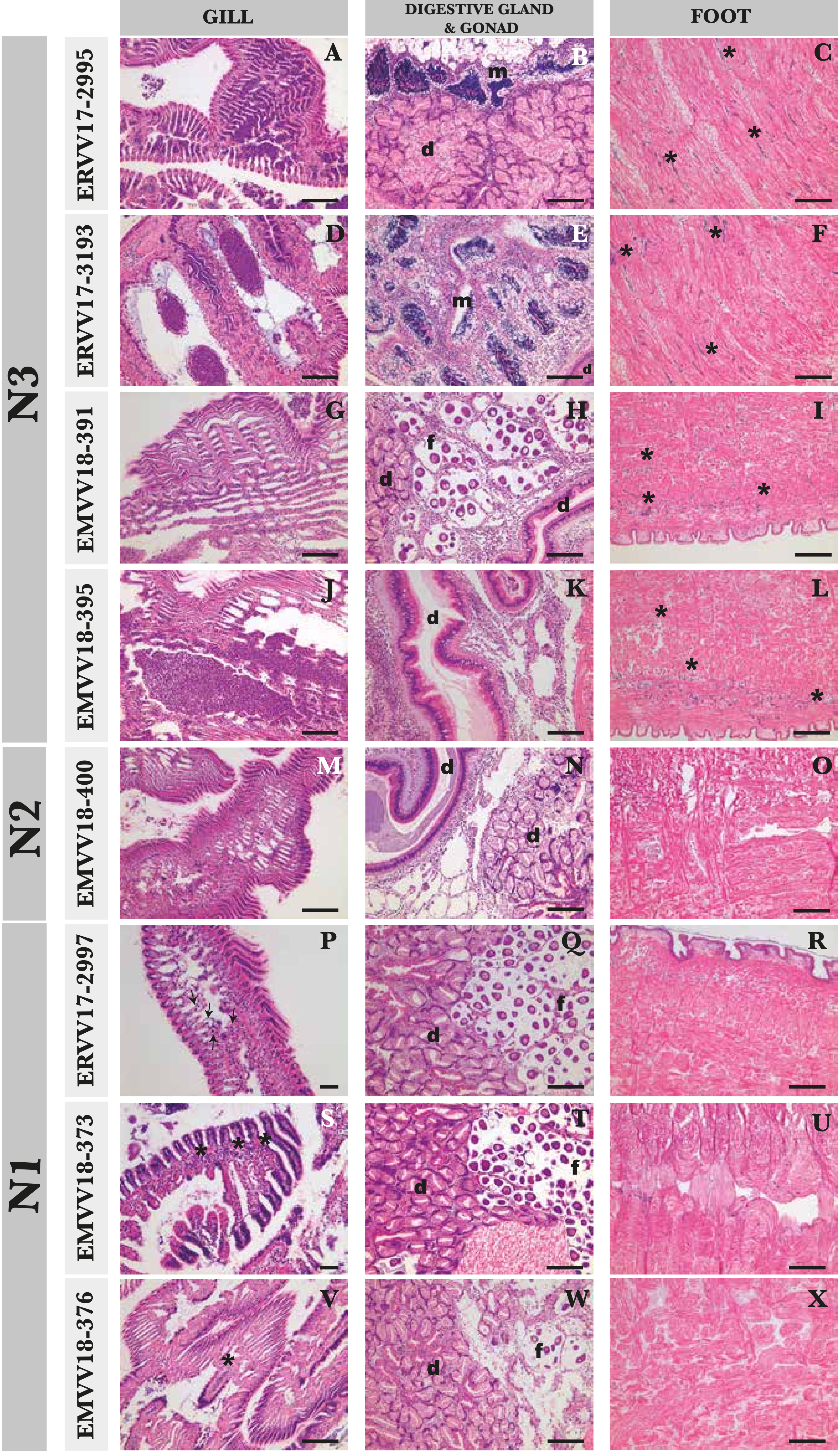

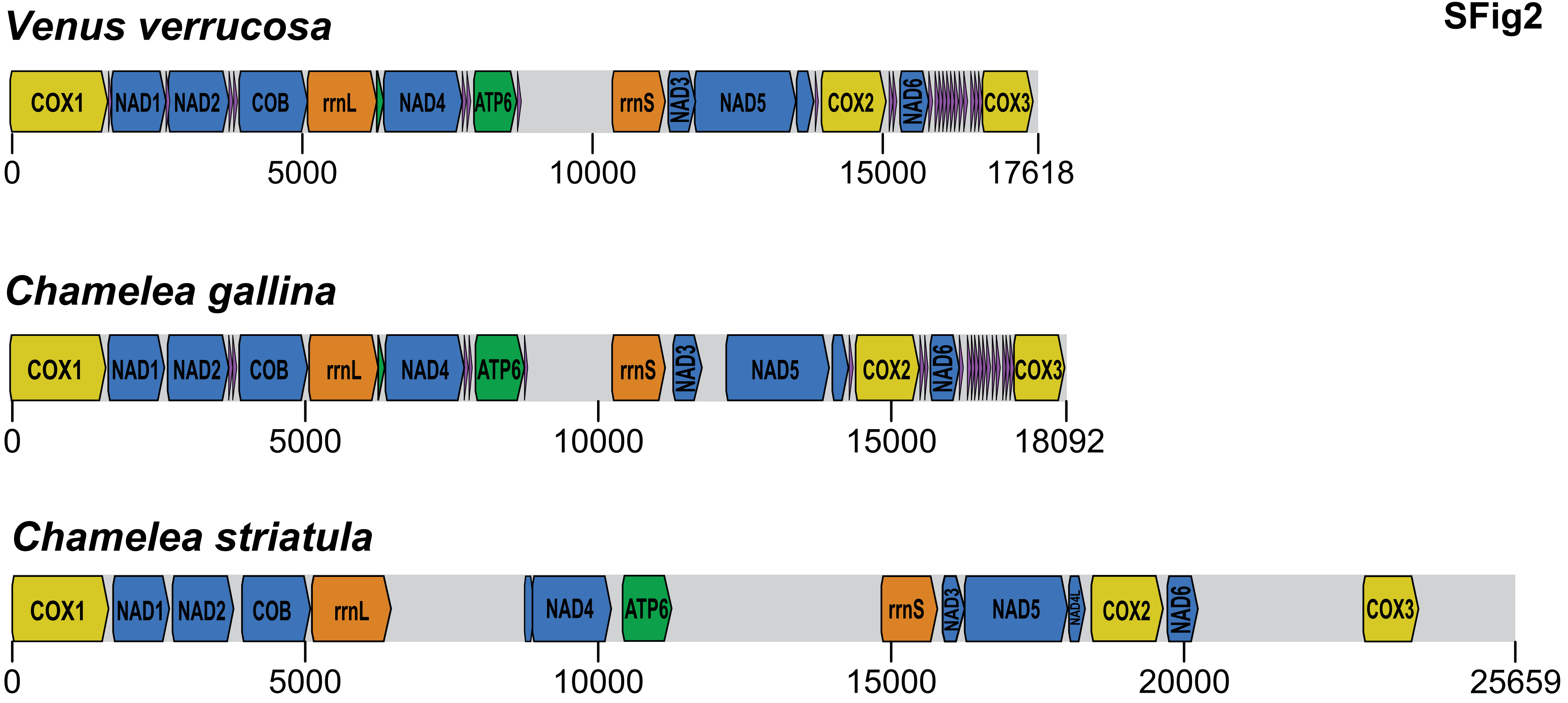
